# New genotypic adaptability and stability analyses using Legendre polynomials and genotype-ideotype distances

**DOI:** 10.1101/2020.05.01.072090

**Authors:** Michel Henriques de Souza, José Domingos Pereira Júnior, Skarlet De Marco Steckling, Jussara Mencalha, Fabíola dos Santos Dias, João Romero do Amaral Santos de Carvalho Rocha, Pedro Crescêncio Souza Carneiro, José Eustáquio de Souza Carneiro

**Affiliations:** Departamento de Agronomia, Universidade Federal de Viçosa, Viçosa, CEP: 36570-900, Minas Gerais, Brazil; Departamento de Biologia Geral, Universidade Federal de Viçosa, Viçosa, CEP: 36570-900, Minas Gerais, Brazil

**Author notes:** Corresponding author –.

## Abstract

Developing cultivars with superior performance in different cultivation environments is one of the main challenges of breeding programs. The current adaptability and stability analyses have limitations, especially when used with trials with genetic or statistical imbalances, heterogeneity of residual variances, and genetic covariance. Thus, adaptability and stability analyses based on mixed model approaches are an effective alternative in such cases. We propose a new methodology for genotypic adaptability and stability analyses, based on Legendre polynomials and genotype-ideotype distances aiming at greater precision when recommending cultivars. We applied the methodology to a set of common bean cultivars throughout a multi-environment trial. We used a set of 13 trials, where they were classified in unfavorable or favorable environments, depending on the average of the cultivars in these trials. The results showed that the methodology allows to predict the genotypic values of cultivars in environments where they were not evaluated with high accuracy values, therefore circumventing the imbalance of the experiments. From these values, it was possible to measure the genotypic adaptability according to ideotypes. The stability of the cultivars was quantified as the invariance of their behavior throughout the trials. The use of ideotypes based on real data allowed a better comparison of the performance of cultivars across environments.

## Introduction

The main objective of a crop breeding program, is to develop cultivars that can replace those that are currently available [1]. In the final stages of a breeding program, the most promising lines are evaluated in trials conducted in different environments, such as different years, places, and seasons. In Brazil, these tests are called Valor de Cultivo e Uso (VCU), and their results are the basis for the cultivar recommendation [2].

Adaptability and stability studies are used to quantify the performances of the genotypes to make recommendations [3]. Adaptability is defined as the ability of a genotype to respond advantageously to its environment, while its stability is related to the predictability of its behavior [4,5]. It is thus possible to identify genotypes that have wide or specific adaptability to favorable or unfavorable environments. Finlay and Wilkinson [4] defined favorable and unfavorable environments as those that result in the average performance of the genotype being above or below the average of all the trials, respectively.

Genotypes that have specific adaptability to favorable environments, have genes that enable them to respond to improved environmental conditions, and should be recommended to farmers who wish to utilize the most current technologies. Genotypes with specific adaptability to unfavorable environments however, may have specific genes that enable them to grow in these environments. These are rustic genotypes and should be recommended to farmers who utilize lower level technologies. In general, rustic genotypes have more genes that tolerate biotic and abiotic stresses, which means they may be favored in more adverse environmental conditions.

In recent decades, several methods to analyze adaptability and stability have been proposed, based on different statistical principles. To identify genotypes that have general or specific adaptability to favorable (requires a high level of technology) or unfavorable (requires a low level of technology) environments, methodologies that are based on linear regression models have shown promise [4–7]. Some of the previous methods to determine the adaptability and stability included the ideotype concept [8–11], and resulted in an improved understanding of the relative behavior of the genotypes from a smaller number of parameters. According to Eeuwijk et al. [12], there are other methodologies to assess the behavior of genotypes that are of note, such as AMMI (Additive Main effects and Multiplicative Interaction) [13] and GGE biplot (Genotype main effects and Genotype × Environment interaction effects) [14]. However, the adaptability and stability analyses still have limitations, especially when used with trials with genetic or statistical imbalances, heterogeneity of residual variances, and genetic covariance. In this context, adaptability and stability analyses that use a mixed model approach are an effective alternative to the traditional analyses [15,16].

Another relevant factor is that traditional methodologies for the analysis of adaptability and stability, consider a priori, that the behavior of a genotype across environments is linear, which may not be true. As a consequence, recommendations based on these methodologies can be biased. This can be outlined by means of reaction norm models via mixed modeling, as they allow for improved modeling of the behavior of the different genotypes, based mostly on orthogonal polynomials. Among this class of polynomials, the Legendre’s polynomials stand out, as they have the ability to describe the structures of variance and covariance between the genetic and environmental components [17].

In this way, the use of the reaction norms obtained from the Legendre polynomials can better quantify the adaptability and stability of a set of genotypes evaluated in different environments, aiming for greater accuracy in cultivar recommendations. Thus, the objectives of this investigation were to propose a new methodology for the analysis of adaptability and genotypic stability, based on Legendre polynomials and genotype-ideotype distances.

## Methods description (step by step)

### Step one: Environmental gradient

The first step is the classification of the trials as an environmental gradient. To define this gradient, trials in which the genotypes are evaluated must be ordered a priori, according to certain classification criteria such as Akaike Information Criterion (AIC) [18], Bayesian Information Criterion (BIC) [19], and Penalizing Adaptively the Likelihood (PAL) [20]. We consequently recommend the index proposed by Finlay and Wilkinson [4], since the adaptability of a genotype is its ability to respond to environmental improvements. The environmental index was determined as follows:

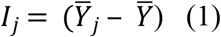

where 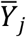 is the average of the genotypes *j-th* trial (*j* = 1, 2, …, *na*, where *na* is the total number of trials) and 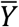 is the general mean. Negative and positive index values indicate unfavorable and favorable trials, respectively.

### Step two: Fitting reaction norm models

Once the environmental gradient is established, different reaction norm models must be adjusted to identify what best quantifies the behavior of the genotypes in the different trials. The number of models to be tested depends on the number of trials used (determines the maximum order of the polynomial), the number of effects included in the model via the Legendre polynomials, and the residual covariance structures.

For the trials conducted in randomized block designs, for example, the model to be adopted was as follows:

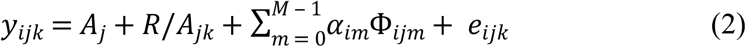

where: *y_ijk_* is the observation of the *i-th* genotype (*i* = 1, 2, … *ng*, where *ng* is the total number of genotypes), in the *j-th* trial (*j* = 1, 2, …, *na*, where *na* is the total number of trials), in the *k-th* block (*k* = 1, 2, 3); *A_j_* is the effect of the trial; *R/A_jk_* is the fixed effect of the blocks within each trial; α_*im*_ is the reaction norm coefficient for the Legendre polynomial of order *m* for the genotypic effects of the genotypes; *Φ_ijm_* is Legendre’s *m-th* polynomial for the *j-th* trial, standardized from −1 to +1 for the *i-th* genotype; *M* is the order of adjustment of the Legendre polynomial for genotypic effects; and *e_ijk_* is the residual random effect associated with *y_ijk_*.

In a matrix, the model above is described as: *y* = *Xb* + *Zg* + *e*, where: *y* is the vector of phenotypic data; *b* is the vector of the fixed effects of the combination of blocks × trials added to the general average; *g* is the vector of genetic effects (assumed to be random); and *e* is the residue vector (random). *X* and *Z* represent the incidence matrix for these effects, respectively. It is assumed that: *g* ~ N (0, *Kg* ⊗ *I_ng_*), and *e* ~ N (0, *I_np_* ⊗ ∑), where *I_ng_* and *I_np_* are identity matrices of the order *ng* (*ng* is the total number of genotypes) and n*p* (*np* is the number of genotypes × the number of blocks), respectively. The symbol ⊗ denotes the Kronecker product. *Kg* is the matrix of covariance coefficients for genotypic effect. *∑* represents the matrix of residual variances.

### Step three: Choosing the best fit model

To select the best fit model, criteria the AIC, BIC and PAL were utilized. These criteria are described as follows:

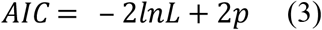

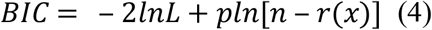

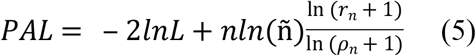

where;

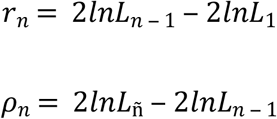

and *lnL* is the logarithm of the likelihood function; *p* is the number of estimated parameters; *n* is the number of observations; *r(x)* is the rank of the fixed effects matrix; and *ñ* is the highest number of parameters for the models.

### Step four: Genetic effects significance

To test the genetic effects, we utilized the Likelihood Ratio Test (LRT) [21], which is as follows:

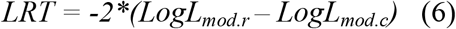

where: *LogL_mod.r_* is the logarithm value of the maximum likelihood function obtained for the reduced model (without the genotypic effect), and *LogL_mod.c_* is the logarithm value of the maximum likelihood function obtained for the complete model.

### Step five: Genotypic values at the original scale

To predict the genotypic values (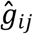 – BLUPs), we suggest the use of the following equation, as proposed by Kirkpatrick et al. (1990):

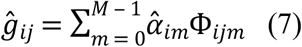

where: 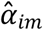 is the reaction norm coefficient of order *m* for the genetic effects of the *i-th* genotype. This equation includes a transformation to the original scale, as using the Legendre polynomials as a covariate affects the scale of the genotypic values.

### Step six: Accuracy at the original scale

The prediction accuracy, also in original scale, is estimated according to the following equation:

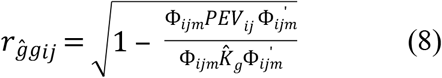

where: 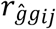 is the correlation between the predicted and real genotype values for genotype *i* in trial *j*, that is, the estimated accuracy; *PEV_ij_* is the Predicted Error Variance of the estimated coefficients for genotype *i* in trial *j*; 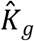 is the covariance matrix of the coefficients, estimated for the genotypic effect.

### Step seven: Genotypic adaptability and stability

To quantify the adaptability and stability of the genotypes, we proposed the use of the genotype-ideotype distance (converted into probability), using four ideotypes: i) genotypes of general adaptability (genotypes of maximum performance in both unfavorable and favorable environments); ii) genotypes of maximum adaptability to unfavorable environments (genotypes of maximum performance in unfavorable environments, regardless of their performance in favorable environments); iii) genotypes of maximum adaptability to favorable environments (genotypes of maximum performance in favorable environments, regardless of their performance in unfavorable environments); and iv) genotypes with minimal adaptability.

From the genotypic values, the adaptability and stability of the genotypes were obtained, according to the estimator below:

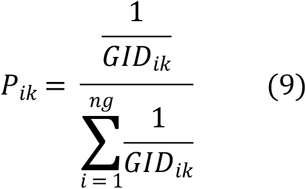

*P_ik_* are the probabilities referring to genotype *i* with regard to ideotype *k*(*k* = 1, 2, 3, 4; where 1 = genotypes of general adaptability; 2 = genotypes of maximum adaptability to unfavorable environments; 3 = genotypes of maximum adaptability to favorable environments; and 4 = genotypes of minimal adaptability); and *ng* is the total number of genotypes. *GID_ik_* is the standardized average Euclidean distance for genotype *i* in ideotype *k*, as given by:

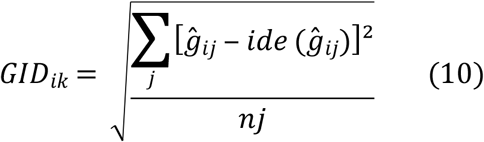

where, if *k* = 1, *j* = 1, …, *na*; if *k* = 2, *j* = 1, …, *nd*; if *k* = 3, *j* = 1, …, *nf*; if *k* = 4, *j* = 1, …, *na*; and *na* is the highest assumed value for *j*; *nd* and *nf* represent the number of unfavorable and favorable environments, respectively; 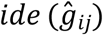 is the ideotype drawn from the standardized genotypic values.

It is important to emphasize that the estimators proposed above also considered the stability of the genotypes’ behavior in relation to the ideotype, through the invariance in multi-environment trials (MET).

We recommend evaluating the performance only in those genotypes that present an accuracy value of at least 80 % in the trials, since the accuracy is indicative of the precision in the prediction of genotypic values. Thus, the average accuracy of the trials considered in the cultivar recommendation will also show values equal to or greater than 80 %. The standard value is based on that of Resende and Duarte [23], who claimed to have at least 80 % accuracy values in VCU trials, when assessing the values of their cultivars.

## Application of the method with *Phaseolus vulgaris L*

### Genetic material

We evaluated 105 common bean cultivars (*Phaseolus vulgaris L*.), 56 of which were Carioca grains and 49 were Black grains. These are the cultivars that have been recommended in Brazil by breeding programs since 1959. The cultivars used, as well as the institutions of origin and year of recommendation, are listed in S1 and S2 tables (supporting information).

### Trials

The trials were conducted in different environments (seasons, years, and places), during the dry and winter seasons, between 2013 and 2018, at the Experimental Stations in Coimbra county – Minas Gerais (Unidade de Ensino, Pesquisa e Extensão – UEPE Coimbra: latitude 20°49’44” S, longitude 42°45’56” W and altitude of 713 meters) and Viçosa – Minas Gerais (Aeroporto, latitude 20°44’38” S, longitude 42°50’40” W and altitude of 654 meters; Horta Nova: latitude 20°45’47” S, longitude 42°49’25” W and altitude of 664 meters; Vale da Agronomia: latitude 20°46’04” S, longitude 42°52’11” W and altitude of 662 meters), thus each MET consisted of 13 trials. Over the years in which the trials were carried out, the cultivars that were recently launched by the breeding programs were included, thus causing a genetic imbalance (variation in the number of cultivars in the trials). The 13 trials and their characteristics are listed in S3 table (supporting information).

The trials were designed in randomized blocks with three replications. The plots consisted of four lines of two meters (m), spaced 0.5 m apart. The treatments used were in accordance with the recommendations for common bean cultures [24]. The evaluated characteristic was grain yield, and they were harvested from the two central lines of each plot. The data were corrected to 13 % humidity and converted to kg ha^−1^.

### Data analysis

To create and organize the environmental gradient, the 13 trials were classified as favorable or unfavorable, according to the environmental index (Eq. 1). We adjusted 14 reaction norm models to identify the model that best quantifies the behavior of the cultivars for grain yield in the MET, with trials ordered according to the environmental index. Among these models, seven were tested considering the homogeneous residual variance and the other seven with heterogeneous diagonal residual variance. The models were adjusted with Legendre’s polynomials, considering the various adjustment orders and based on the general model presented in Eq. 2.

Different degrees of orthogonal Legendre polynomials were fitted to determine the best model (lowest mean square error and greater parsimony). The reaction norm models were compared using the AIC (Eq. 3), BIC (Eq. 4), and PAL (Eq. 5) criteria. The LRT test, presented in Eq. 6, was used to test the significance of the genetic effects. The genotypic values for each cultivar (BLUP), in each trial, were predicted according to Eq. 7. Prediction accuracy was estimated according to Eq. 8.

By using the BLUPs, the adaptability and stability of the cultivars were determined, aiming at the recommendations of the cultivars. In this way, we have calculated the probabilities for the recommendations of the cultivars, using the distance of the genotypes in the functions of the ideotypes. The probability values were obtained using Eq. 9.

To view the results, the ten cultivars with the highest probability were selected to plot their curves with their respective reaction norms, for the three ideotypes, as we chose not to include ideotype IV, since it makes no sense to recommend cultivars of minimal adaptability. The BLUP of each cultivar was added, plus the environment average, and the general average, as well as two witnesses, Pérola (Carioca bean) and Ouro Negro (Black bean), for comparison purposes. These two cultivars were selected as witnesses, as they are used as references for the productivity and quality of grain in consumer markets for the Carioca and Black beans, respectively [25]. The accuracy values (S4 table) and recommendation probability values (S5 table) are available in the supporting information.

### Software used

The joint analysis was carried out using ASREML software [26]. The study of the adaptability and stability of cultivars was carried out using R [27]. The code for the analyses is available in the S1 code.

## Results

The environmental index values, according to Finlay and Wilkinson [4], are shown in table 1. Positive index values indicate favorable environments, while negative values indicate unfavorable ones [3]. Trials 12, 9, 4, 10, 8, and 6 were classified as unfavorable environments, while trials 5, 2, 3, 7, 1, 11, and 13 were favorable

**Table 1:** Trials evaluated with their environmental index.

| **Trial** | **Description** | **Environmental gradient** |
| --- | --- | --- |
| 12 | Dry/2017/Aeroporto | -1028.89 |
| 9 | Dry/2016/UEPE Coimbra | -868.67 |
| 4 | Winter/2013/Vale da Agronomia | -607.25 |
| 10 | Winter/2016/UEPE Coimbra | -500.76 |
| 8 | Dry/2016/Aeroporto | -466.43 |
| 6 | Winter/2015/UEPE Coimbra | -167.35 |
| 5 | Dry/2015/UEPE Coimbra | 31.02 |
| 2 | Dry/2013/Vale da Agronomia | 106.80 |
| 3 | Winter/2013/Coimbra | 127.71 |
| 7 | Dry/2016/UEPE Coimbra | 259.39 |
| 1 | Dry/2013/Coimbra | 486.60 |
| 11 | Winter/2016/Horta Nova | 758.98 |
| 13 | Winter/2017/UEPE Coimbra | 1868.83 |

We found that the different criteria (AIC, BIC, and PAL) pointed to different models as having a better fit. The AIC criterion identified model Leg.6.D, which has a diagonal structure for the residues and a grade six for the Legendre polynomials, as having the best fit (Table 2). The BIC and PAL criteria however, identified the Leg.5.D model as having the best fit. The AIC and BIC criteria prioritize, respectively, efficiency and consistency in their choices of model [28,29]. Corrales et al. [29], using simulated data, reported that when the true model was among the candidate models, the PAL and BIC criteria selected the same model. Furthermore, when the PAL and AIC criteria were used, the model selection was not always the same. When the real model was unknown, the AIC was more precise in choosing the best model, compared to the BIC. According to Vrieze [30], for very complex models (which include a high number of parameters) the BIC criterion was preferred over the AIC. Corrales et al. [29] stated that the PAL criterion simultaneously considers the consistency and efficiency of a model and should, therefore, be preferred over the AIC and BIC criteria when choosing models. The model ultimately chosen was Leg.5.D.

**Table 2.** Different fitted models using the Legendre polynomials (Leg).

| Model<sup>1</sup> | DEG | p | LOG L | AIC | BIC | PAL | LRT |
| --- | --- | --- | --- | --- | --- | --- | --- |
| Leg.0.H | 0 | 2 | Converged | 11307 | 11319 | 11303 | 377.68 |
| Leg.1.H | 1 | 4 | Converged | 11288 | 11312 | 11280 | 400.6 |
| Leg.2.H | 2 | 7 | Converged | 11255 | 11299 | 11257 | 438.8 |
| Leg.3.H | 3 | 11 | Converged | 11218 | 11286 | 11229 | 484.1 |
| Leg.4.H | 4 | 16 | Converged | 11191 | 11291 | 11217 | 520.7 |
| Leg.5.H | 5 | 22 | Converged | 11150 | 11286 | 11197 | 574.5 |
| Leg.6.H | 6 | 29 | Converged | 11134 | 11315 | 11243 | 603.7 |
| Leg.0.D | 0 | 14 | Converged | 11007 | 11094 | 10979 | 507.6 |
| Leg.1.D | 1 | 16 | Converged | 10983 | 11083 | 10951 | 534.9 |
| Leg.2.D | 2 | 19 | Converged | 10928 | 11046 | 10930 | 595.9 |
| Leg.3.D | 3 | 23 | Converged | 10865 | 11008 | 10885 | 667.2 |
| Leg.4.D | 4 | 28 | Converged | 10740 | 10914 | 10779 | 802.7 |
| Leg.5.D | 5 | 34 | Converged | 10648 | 10859 | 10730 | 906.2 |
| Leg.6.D | 6 | 41 | Converged | 10645 | 10900 | 10881 | 922.9 |

Model structure, degree polynomial for the genetic effect (DEG), number of parameters (p), LOG L convergence, Akaike information criterion (AIC), Schwarz Bayesian information criterion (BIC), Penalizing adaptively the likelihood (PAL) and Likelihood Ratio Test (LRT).
<sup>1</sup>The tested models can assume homogeneous (H) or diagonal (D) residual variance structure.

Based on the chosen model (Leg.5.D), the random effects of the cultivars were modeled as linear functions using the Legendre polynomials, with grade five and heterogeneous residual variance (diagonal). This resulted in 34 estimated parameters, 13 of which were associated with residues, that is, one for each trial, and 21 related to the model’s genotypic components. It is of note that the genetic effect was significant with the LRT test for all fitted models, indicating high variability between the cultivars evaluated (Table 2).

The average accuracy for the prediction of the BLUPs for each cultivar, based on the Leg.5.D model, are shown in Fig 1. We found that the average accuracy of predictions was greater when more trials were used to evaluate the cultivars. The accuracy observed for the cultivars that were present in the 13 environments was the highest, while the accuracy estimates for the cultivars evaluated in only two environments were the lowest. It can also be seen in figure 1, that in trials six and eight (ordered according to the environmental gradient), the accuracy estimates were relatively low. It is also noteworthy that the estimates of genotypic variance in these two trials were also lower than in the others (data not shown). The accuracy values of each cultivar in each environment are available in S4 table (supporting information).

**Fig 1:**
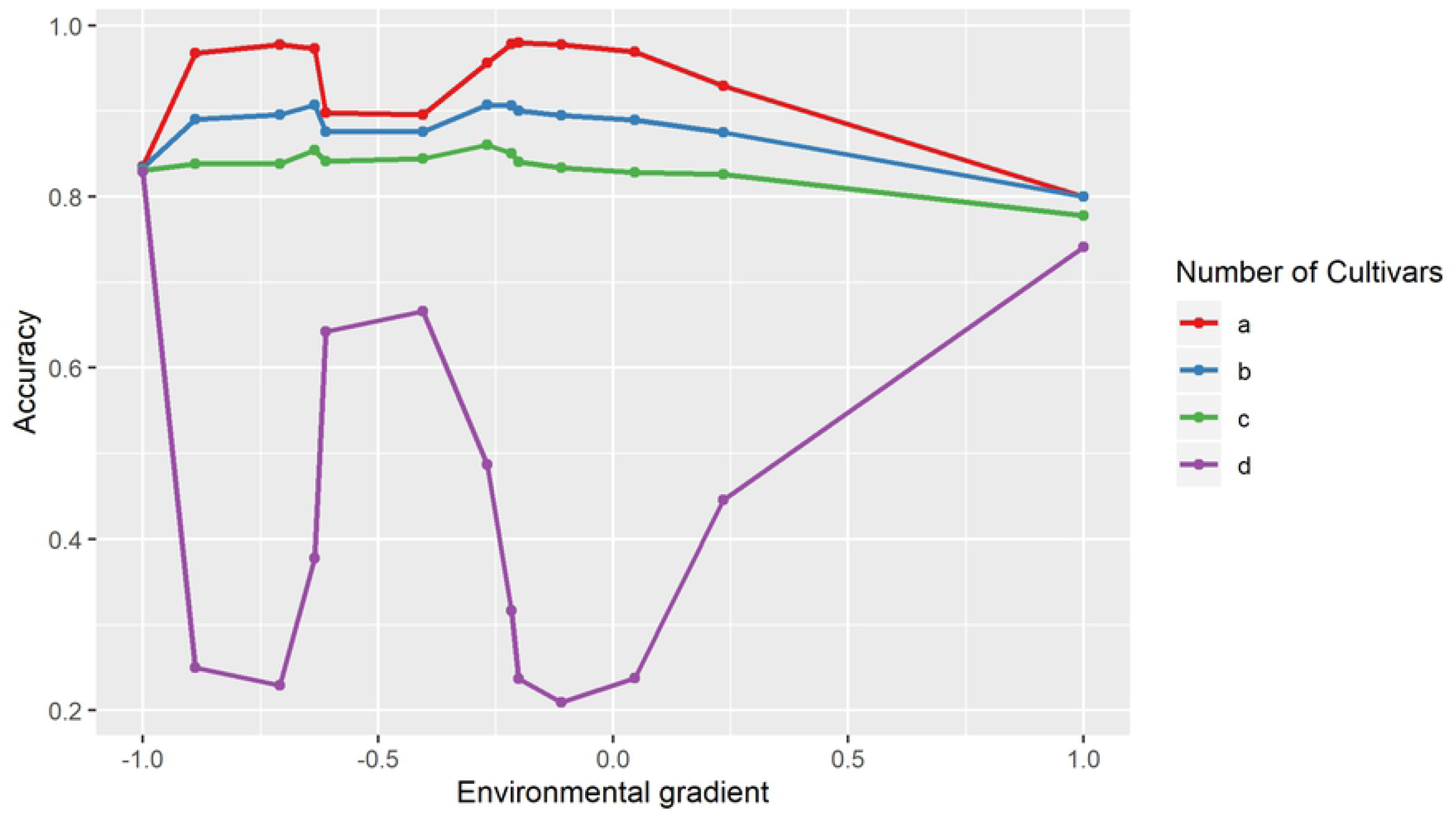
Average accuracy of the prediction in each trial for the genotypic values of the cultivars. a) Cultivars evaluated in 13 trials (80 cultivars); b) cultivars evaluated in nine trials (20 cultivars); c) cultivars evaluated in six trials (four cultivars); and d) cultivars evaluated in only two trials (one cultivar). The trials are ordered according to the environmental index (Table 1).

Using the proposed reaction norm methodology, the adaptability and stability of 100 of the 105 cultivars was quantified. These 100 cultivars were evaluated in at least nine of the 13 trials, with the accuracy in predicting their genotypic values, equal to or greater than 80 %, including for those trials in which the cultivars were not evaluated (S4 table).

According to Eq. 9, the cultivars were recommended by comparing them with the four proposed ideotypes (four scenarios): cultivars of general adaptability, cultivars of maximum adaptability to unfavorable environments, cultivars of maximum adaptability to favorable environments, and cultivars of minimal adaptability. The probability values of each cultivar in each scenario are presented in S5 table.

Fig 2 shows the reaction norm curves of the ten bean cultivars with the highest potential (highest probability value), considering the general adaptability scenario (ideotype – maximum performance genotypes in both unfavorable and favorable environments), as well as the cultivars used as witnesses (Pérola and Ouro Negro). The probability of each cultivar was calculated according to eq. 9, in relation to the ideotype for the scenario of general adaptability. Among the ten selected cultivars, six had the Carioca grain type (BRS Estilo, IAC Formoso, IAC Imperador, IPR Andorinha, IPR Campos Gerais and VC 15), and four had the Black grain type (BRS Agreste, IPR Tiziu, IPR Tuiuiú and VP 22). The IPR Campos Gerais cultivar surpassed the Pérola cultivar in all trials, while the VP 22 cultivar surpassed the Ouro Negro cultivar in all trials.

**Fig 2:**
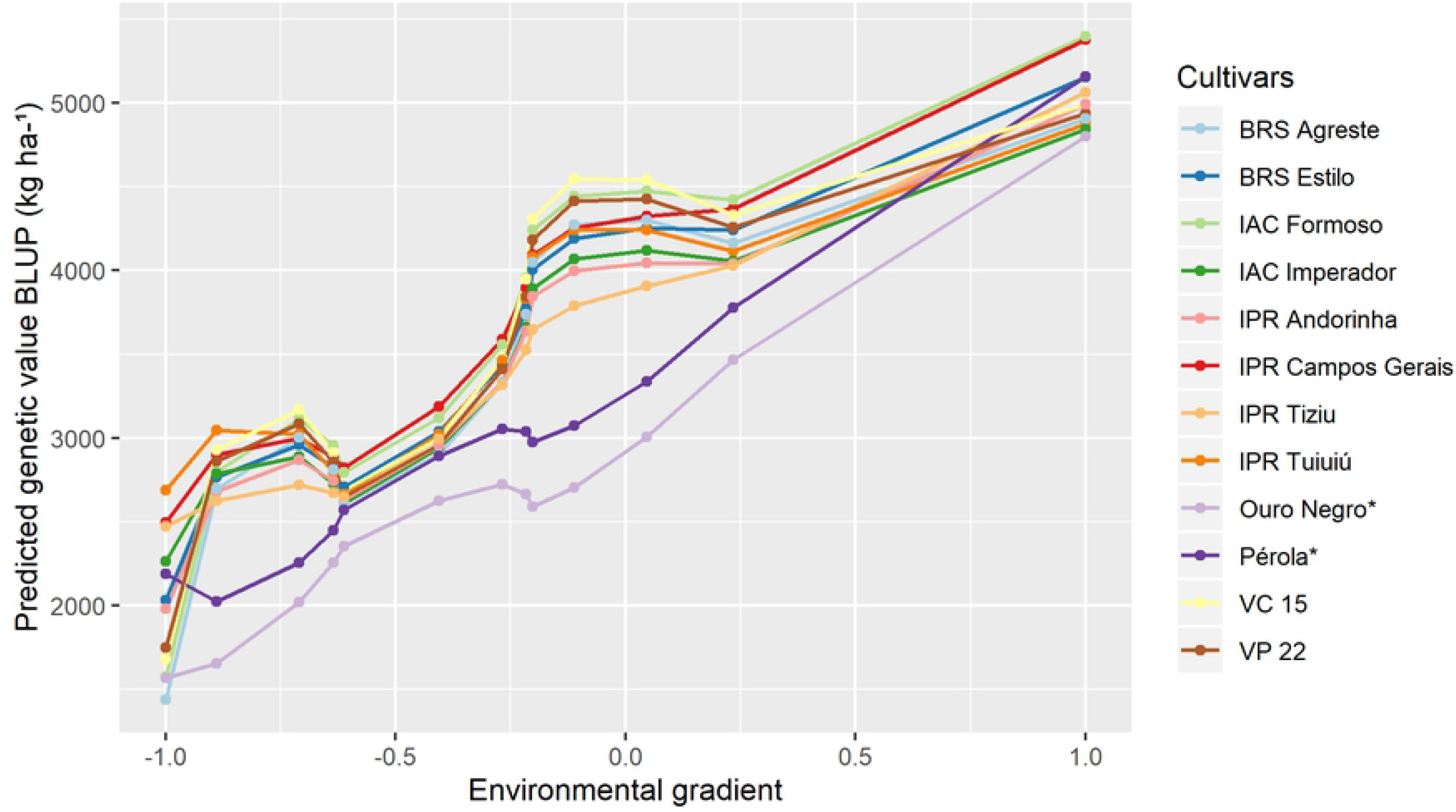
Cultivars of Carioca and Black bean of general adaptability according to the ideotype. The trials are ordered according to the environmental index (Table 1). *Cultivars used as witnesses.

The reaction norm curves of the ten bean cultivars with the greatest potential (highest probability value), considering the scenario of maximum adaptability to unfavorable environments (ideotype – maximum performance genotypes in unfavorable environments, regardless of their performance in favorable environments), as well as the cultivars used as witnesses, are presented in Fig 3. Of the ten selected cultivars, seven had Carioca grain (BRS Estilo, IAC Formoso, IAC Imperador, IPR Andorinha, IPR Campos Gerais, IPR Tangará and VC 15) and three had Black grain (IPR Tiziu, IPR Tuiuiú and VP 22). The cultivar IPR Campos Gerais surpassed the cultivar Pérola in all trials, and the IPR Tuiuiú, IPR Tiziu, and VP 22 cultivars exceeded the Ouro Negro cultivar.

**Fig 3:**
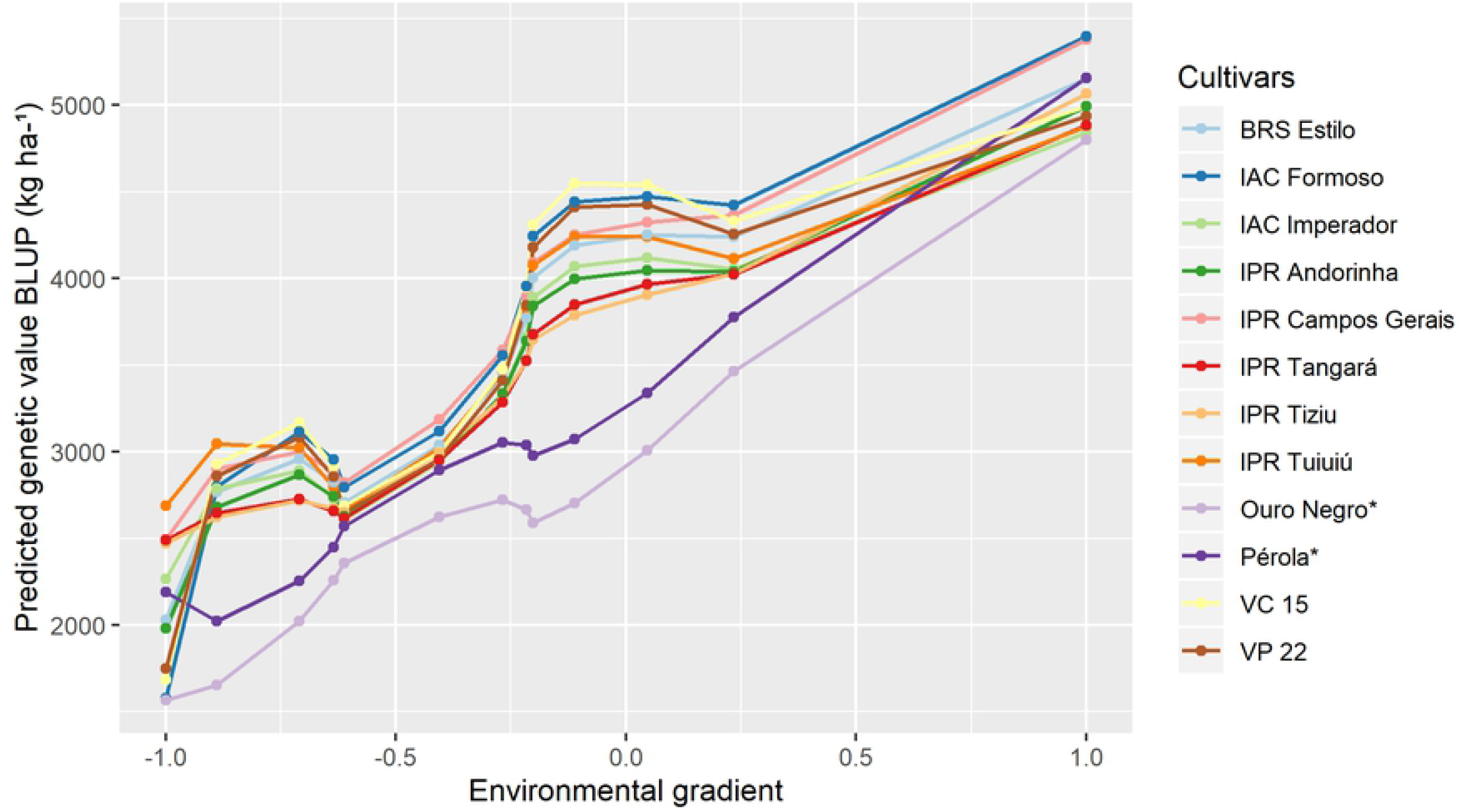
Cultivars of Carioca and Black bean of maximum adaptability for unfavorable environments according to the ideotype. The trials are ordered according to the environmental index (Table 1). *Cultivars used as witnesses.

In Fig 4, the reaction norm curves for the ten cultivars with the highest potential (highest probability value), considering the scenario of maximum adaptability to favorable environments (ideotype – maximum performance genotypes in favorable environments, regardless of their performance in unfavorable environments), as well as the cultivars used as a witness, are shown. Of the ten selected cultivars, seven had Carioca grain (BRS Estilo, IAC Formoso, IAC Imperador, IPR Andorinha, IPR Campos Gerais, IPR 139 and VC 15) and three had Black grain (IPR Agreste, IPR Tuiuiú and VP 22). The IPR Campos Gerais cultivar surpassed the Pérola cultivar, in all trials, and the IPR Agreste, IPR Tuiuiú, and VP 22 black bean cultivars exceeded the Ouro Negro cultivar, in all trials.

**Fig 4:**
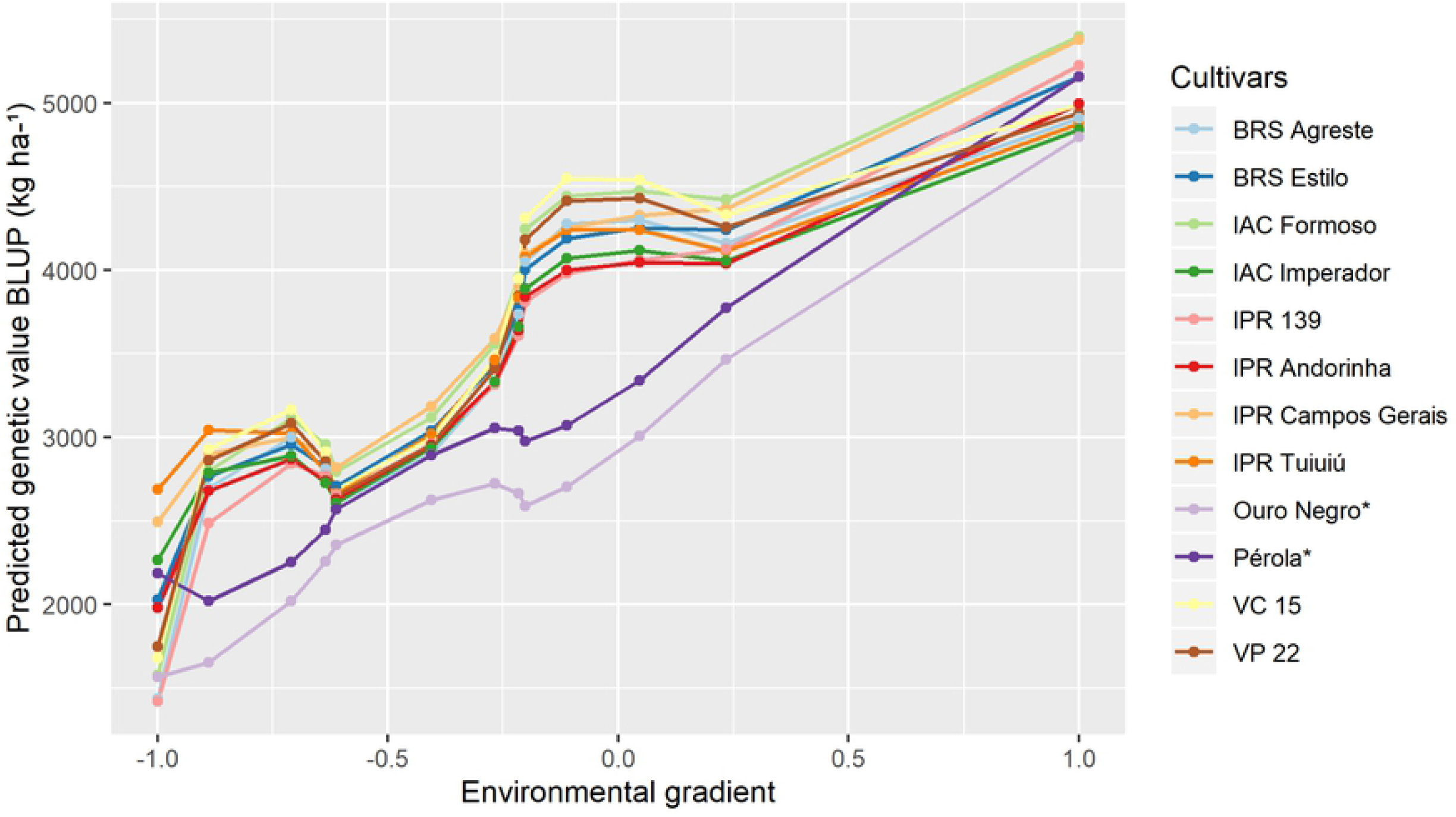
Cultivars of Carioca and Black bean of maximum adaptability for favorable environments according to the ideotype. The trials are ordered according to the environmental index (Table 1). *Cultivars used as witnesses.

## Discussion

Rating the variations of a set of trials, according to an environmental gradient, is essential when using methods based on linear regression that aim to quantify the adaptability of a cultivar. Finlay and Wilkinson [4] proposed using the average performances of the cultivars in each trial as a gradient, and estimating an environmental index using the differences between the average of the cultivars evaluated in each trial and the general average of the cultivars in all trials. Additionally, the fit of the regression model for each cultivar was made according to its performance, relative to the environmental index, in order to increase the values. The lack of an environmental gradient complicates the interpretation of the behavior of the genotypes in the face of the environmental variations [4].

When classifying the trials with the environmental index (in favorable or unfavorable environments), it was observed that the seasons, places, and years in which the trials were conducted did not determine the classification, as the trials from the same place and year could have very different results (trials 7 and 9), while those from different seasons, places, and years could be very similar (for example, environments 1 and 11). It should be noted that trial 9 was planted 44 days after trial 7, which may be one of the justifications for the different environmental index values. These results could be caused by edaphoclimatic variations, as well as variations in the incidence of pests and diseases in the environments in which the cultivars were evaluated, resulting in genotype by environment interactions (GEI). Several authors have also previously [31–34] reported the influence of these factors on the environmental classification. For Ramalho et al. [35], the most significant contributions to the GEI in the bean cultures were due to the combinations of cultivar × season and cultivar × years.

The development of methods to model GEI is coupled with the availability of more genotypic and environmental information, in line with the advances in data collection and analysis. The first analyses were based on analysis of variance [36,37], with a single parameter to interpret the adaptability and stability. The advances with the development of new methodologies however, are based on regression analysis, with interpretations based on more parameters, such as the average, the regression coefficient, the regression deviation, and new definitions of adaptability and stability [4,7,38].

Currently, the effects of genotypes and environmental conditions can be modeled by phenotypic values in regression with genetic markers and in environmental covariates, via mixed models [39]. However, these models based on regression, consider a priori that the genotype behavior is predetermined, based on linear regression equations, which may not equate to the genotypes actual behavior. Thus, reaction norm models in conjunction with Legendre polynomials are used to establish the order of the polynomials of the regression parameters later, according to the behavior of the genotypes in a series of environments (in a MET). Additionally, the mixed model approach also allows for the genotypic values of individuals to be predicted, as adaptability and stability are genotypic, and not phenotypic.

The use of reaction norm models associated with the use of orthogonal polynomials has been used mainly in animal breeding, defined as random regression, where the behavior of the genotypes over a period of time is described, mainly using covariance function information [40–42]. However, there are only a few studies in the literature that use reaction norm models with plants.

According to Ni et al. [43], reaction norm models allow for the adjustment of an individual’s genetic effects with their exposure to the environmental effects, so that the genotypes are adjusted as a nonlinear function of a continuous environmental gradient. The adjustment of reaction norm models, as a function of the environmental gradient, considering Legendre polynomials, captures more adequately the behavior of the genotypes in a MET. The fact that an individuals’ behavior is not predetermined, is an advantage of the proposed methodology in relation to the traditional methods of analysis of adaptability and stability.

To quantify the adaptability and stability using reaction norm models, the prediction accuracy represents the reliability in the evaluation of the behavior of the evaluated genotypes in different environments. In this work, most of the accuracy estimates obtained for each cultivar in each environment were greater than 80 %, which also resulted in an average accuracy of the 13 trials that was higher than this value. In the VCU trials, Resende and Duarte [23] recommended that the accuracy should be at least 80 %. Other previous investigations have also highlight the importance of prediction accuracies, using the reaction norm models in plant breeding experiments [39,44,45].

Another advantage of the proposed methodology, using reaction norm models, is the prediction of genotypic values for the cultivars for environments in which they were not evaluated, when the MET presents genetic imbalance. When using experiments with unbalanced data, or just a sample of the cultivars, the prediction accuracy estimates tend to be lower, and the model may not be efficient in evaluating the performance of the cultivars [46,47]. Cargnelutti Filho and Storck [48], affirmed that the accuracy has a direct relationship with the genotypic variance, and an inverse relationship with the residual variance. In this investigation, we observed accuracy estimates of at least 80 %, when the cultivars were evaluated in at least 9 of the 13 environments (Figure 1).

For Smith et al. [49], using accurate information for the behavior of the cultivars, allowed breeders to choose the best varieties, according to the needs of farmers, in order to maximize profitability and food security. One of the difficulties in assessing the behavior of a group of cultivars over MET was due to the fact that new genotypes were included in the trials over the years, in addition to the loss of information due to problems that occurred over the trials, resulting in genetic and statistical imbalances. In this context, Resende [16,50] states that the mixed model approach is a better alternative, for the analysis of such trials.

As noted, only 12 cultivars of superior performance were found in Fig 2–4, with eight carioca bean cultivars and four black bean cultivars, instead of 30 cultivars (10 per figure). This was because there were some cultivars that were widely adaptable and highly stable that were selected for more than one scenario, such as the IPR Campos Gerais and IPR Tuiuiú.

Cultivars with high phenotypic averages for high yield were identified, but they were not included in figures 2, 3, and 4, as those selected by the reaction norm models. This can be explained by the fact that the methodology when calculating the probability of each cultivar that was based on the cultivar-ideotype distance penalizes cultivars that showed great variation in their productivity during the trials, even if they presented high general averages. Thus, the reaction norm models can also quantify the stability of cultivars, defined as the variation across environments. Eeuwijk et al. and Van Oijen and Höglind [12,51] also reported this property of reaction norms. It is also worth mentioning that the use of the ideotype that was established from the data itself, had the advantage of comparing the genotypes with a real situation observed for that MET, since the ideotype is defined as the maximum value predicted in each trial.

The reaction norms, based on the mixed models, can also model the heterogeneity of the genetic variations and correlations between the environments, in addition to the spatial trends in the trials [22]. Furthermore, these models allow for more accurate estimations of the genotypes in the trials, as well as better estimations of the genetic parameters, such as heritability, variances, covariances, and genetic correlations, while they become more difficult in models with only fixed effects [12].

VCU tests are the basis for the recommendation of a cultivar and it is required that they are carried out in various locations, seasons, and years of the macro-region where the cultivar is being recommended. Thus, the recommended cultivars are those with higher general averages across the environments, that is, wide adaptability. Cultivars with these behavior, beyond the interests of the breeders making the recommendations, are also of interest to the farmers, as beans are mostly cultivated by small farmers, who buy grains from other producers and regions to use as seed. Thus, there is often an overlap and lack of control as to what the planting season and region actually are for a cultivar, and its official recommendations [52].

It is expected that cultivars with maximum adaptability to unfavorable environments will be more desirable for these unstructured conditions. These environments can be described as having low levels of technological investment, which can be normal in small-scale agriculture [3]. In addition, adverse conditions caused by the climate, such as a lack or excess of rain and incidence of pests and diseases also contribute to the characterization of environments as unfavorable. Thus, it is desirable that cultivars that are recommended for unfavorable environments maintain a satisfactory standard of productivity, even in stressful situations, whether this is due to a lack or excess of any factor. However, in a situation of improvement of the environment, these cultivars will not be responsive to this increment of environmental quality. This illustrates the definition of adaptability as presented by Cruz et al. [3], as the differential response of cultivars due to a stimulus from the environment.

However, for the cultivars that are identified for favorable environments, we see the opposite behavior. It is expected that cultivars adapted to these locations would normally respond satisfactorily to environmental improvements, reaching high levels of yield in order to return the investment made, since these environments have high technological use, such as irrigation and precision agriculture, and are commonly run by large scale practices. However, with inferior conditions, such as climatic adversity, they tend to have low production [3]. It was observed that the strains VP 22 and VP 33 showed superior performance in environments classified as favorable. The fact that the experiments conducted by the Programa Feijão – UFV, who are responsible for the selection of these strains and always utilize optimal cultivation conditions (fertilization, irrigation, and pest and disease control), may explain this.

The maintenance of productivity in different environments is explained by the response to the environmental stimulus, being caused by the differential expression of the genes present in each individual. In this way, the adaptability and stability indicated in the reaction norm curves of the cultivars, provides information regarding their capacity to express phenotypes that may better adjust to the environmental conditions [53]. In this sense, one way to improve the adaptability of cultivars to the different environments in which they will be cultivated, is to pyramid the genes of maximum expression in both the unfavorable and favorable environments. The superior cultivars in each studied scenario were developed in different breeding programs from four institutions (EMBRAPA, UFV, IAC, and IAPAR). This is indicative of the effort and success of these breeding programs, as well as the genetic diversity between them, since the breeding programs are independent, with their own parental lines. Possobom [54] demonstrated that cultivars originating from the same institution are usually more related, while cultivars from different institutions belong to different groups of dissimilarity. Thus, these outstanding cultivars also have the potential to be used in bean improvement programs.

## Conclusion

The reaction norm methodology to evaluate the adaptability and stability of cultivars appears to be an alternative in the evaluation of multi-environment trials, since it enables genetic and statistical imbalances to be addressed, as well an improved evaluation of cultivar behavior.

## Acknowledgments

We would like to thank the students of the Programa Feijão for their contribution with the help of data collection for this work. We would like to thank the CNPq (Conselho Nacional de Desenvolvimento Científico e Tecnológico), FAPEMIG (Fundação de Amparo à Pesquisa do Estado de Minas Gerais) and CAPES (Coordenação de Aperfeiçoamento de Pessoal de Nível Superior) agencies for their financial support. We would like to thank Editage (www.editage.com) for English language editing.

## Supporting information

**S1 Table. Carioca bean cultivars, institutions of origin and year of recommendation.**

**S2 Table: Black bean cultivars, institutions of origin and year of recommendation.**

**S3 Table: Description of the trials.**

**S4 Table: Accuracy of 105 cultivars in each trial.**

**S5 Table: Recommendation probability values for each cultivar in each scenario.**

**S1 Code: Script for analyses.**

